# Glial Cell Responses to constant low Light exposure in Rat Retina

**DOI:** 10.1101/2021.05.10.443423

**Authors:** Manuel Gastón Bruera, María Mercedes Benedetto, Mario E Guido, Alicia Laura Degano, María A. Contin

## Abstract

Retinal damage promoted by constant illumination of low intensity resulted in a diminution in classical photoreceptors cells. Glial cells exert profound effects on neurons, vasculature and other glial cells. Macroglia and microglia with specific morphological, physiological, and antigenic characteristics may play an essential role in both the maintenance and control of retinal homeostasis, or to exert mechanisms that promote cell death. The role of glial cells and immune function in the pathogenesis promoted by low light is poorly understood. We performed glial cells characterization along the time-course of retinal degeneration induced by chronic exposure to low intensity of light in Wistar rats. We exposed the animals at constant light from 2 to 8 days and assessed the retinal glia. After 6 days of light exposure, retinas presented increased levels of GFAP, a macroglia marker and microglia markers Iba1 and CD68 displayed increased mRNA levels after 6 days. The number of Iba1 positive cells increased in the outer nuclear layer, showing ameboid morphology with thicker processes characteristic of microglial activated cells. The expression levels of immune mediators TNF-□ and IL-6 were also significantly increased after 6 days. Finally, chemokines analysis showed that CX3CR1 and CCL2 expression levels were significantly elevated after 6 days. Hence, all the events of glial activation occurred after 5-6 days of constant light exposure, when the number of cells of the outer nuclear layer has already decreased significantly. Herein we demonstrated that glial and immune activation are secondary to neurodegeneration; in this scenario, our results suggest that photoreceptor death is an early event that may be induced by phototransduction-dependent mechanisms.

## INTRODUCCION

The effects of excessive artificial light sources called light pollution could have direct consequences on retinal health. Constant exposure to different wavelengths and intensities of light promoted by light pollution may produce retinal degeneration (RD) as a consequence of photoreceptors or retinal pigment epithelium cells death [1]. Due to the wide range of light sources, the retinal damage is usually divided into two classes: (a) damage produced by low irradiance levels of light, mediated by the activation of rhodopsin in photoreceptor cells and (b) damage produced by a brief exposure to high irradiance–(bright light), where injuries occur at short times of exposure [2]. The relationship between photoreceptor degeneration and glia activation in retinal light injury remains poorly understood at the molecular level and it depends on light source, intensity or exposure times; also, differences between *in vivo* / *in vitro* models were found [3–5].

Previously, we demonstrated that constant illumination of low intensity white light (200 lux) using LED devices produced RD in Wistar rats. We observed a significant reduction in the outer nuclear layer (ONL) after 6 days of constant stimulation (LL6) [6]. Moreover, at early exposure times (2 days; LL2), we demonstrated rhodopsin hyper-phosphorylation on serine 334 residue, suggesting that rhodopsin phosphorylation may be affected by prolonged phototransduction mechanism activity, as one of the first events [6]. Also, our results suggest that an apoptosis caspase-3 independent mechanism may be involved, i.e. calpain-dependent mechanism or other pathways such as necroptosis. Interestingly, markers of oxidative stress increased after 5 days of constant light stimulation (LL5) in the ONL and correlated with a reduction in docosahexaenoic acid (DHA) LL6. DHA is the major component of rod outer segment (OS) membrane and it is highly vulnerable to oxidation; thus, oxidative stress processes may affect the OS structures after low light exposure [7]. Therefore, we consider that constant low light-induced photoreceptor cell death in Wistar rats is a useful model for studying the mechanisms involved in phototransduction defects during the early stages, in order to define the main events leading to RD. The retina, as part of the central nervous system (CNS), is nourished by minority cellular subsets, the glia and vasculature. Although glia constitutes a small fraction of the retina by cell number, it exerts profound effects on neurons, vasculature and other glial cells. Retinal glial cells are subdivided into macroglia (Müller cells and astrocytes) and microglia (the resident immune cells) with specific morphological, physiological and antigenic characteristics [8, 9]. Müller cells and astrocytes; Müller cells comprise 90% of the retinal glia and anatomically define the distal and proximal borders of the retina since they spread through the retina from the inner limiting membrane to the distal end to ONL (54). They provide homeostatic and metabolic support to photoreceptors and other neurons required for normal neuronal activity [10]. Müller cells maintain the viability of photoreceptors and neurons; they direct light onto photoreceptors, recycle the retinoids in an alternative visual cycle [50] and provides structural stabilization of the retina, and modulate immune and inflammatory responses [10]. In pathological conditions, a reactive gliosis response is induced, which may have effects aimed to protect the retina against further damage and maintenance of homeostasis [11], or to produce cytotoxic effect, increasing the susceptibility to stressful stimuli [12]. Reactive gliosis has been described in different retinal pathologies, including age-related macular degeneration (AMD), diabetes retinopathy, glaucoma, retinal detachment, or retinitis pigmentosa [10]. Thus, understanding the role of microglia in both, protective and cytotoxic effect, would help treating the retinal physiopathology. Astrocytes are the other type of macroglia in the mammalian retina and play important roles in retinal development and hemodynamics. In response to neuronal activity and neurotransmitter release, astrocytes produce vasoactive compounds and promote vasodilation or constriction of retinal blood vessels [13].

The microglia is a type of retinal glial cells that contributes in the immunity-mediated protective mechanisms, displaying specific morphological, physiological, and antigenic characteristics. Microglial cells act as phagocytes and, together with perivascular cells, form a network of immune effector cells in CNS [14, 15]. In the retina, microglia represent the resident tissue macrophages and play important roles in retinal homeostasis, recovery from injury and progression of disease. Under physiological conditions, microglial cells are located in the inner plexiform layer (IPL) and outer plexiform layer (OPL) participating in neuron-macroglia interaction to maintain the cellular homeostasis. Microglia also fulfill a number of tasks necessary for the physiological functions in the healthy retina; the contribution of microglia in maintaining the purposeful and functional histo-architecture of the adult retina is indispensable, expressing receptors and releasing neuroprotective and anti-inflammatory factors playing a critical role in host defense, immunoregulation, as well as for tissue repair [16]. Moreover, adequate resident microglial population is necessary for proper retinal blood vessel formation [17] and the glial distribution through the retinal layers is well documented as well [18, 19]. Therefore, so-called “resting” microglial cells in the healthy retina play a vital role maintaining the physiological homeostasis of the retina. In pathological conditions, the “activated” microglia denote different functions including migration and morphological changes [17, 20, 21], acquiring an ameba-like shape, the ability to migrate into the damaged region and expressing a number of pro-and anti-inflammatory molecules. The classical photoreceptor ONL, which is devoid of microglial cells in healthy retina, becomes colonized by these cells in conditions that induce massive ONL cell degeneration, such as inherited photoreceptor degeneration [21] or light/laser injury [22, 23]. In some age-related diseases, as AMD, the activated microglia can be neurotoxic and lead to the degeneration of photoreceptors, thereby contributing to typical chronic inflammation [8, 14].

As retinal glial cells play an essential role in the maintenance and control of retinal homeostasis, in the present work we aimed to characterize glial cells response along the time-course of RD induced by chronic exposure to low intensity LED light in Wistar rats. The use of such a model in the comprehensive study of RD may provide a basis for future strategies for preventing or delaying visual cell loss.

## MATERIALS AND METHODS

### Animals

All animal procedures were performed in accordance with the ARVO statement for the use of animals in ophthalmic and vision research, which was approved by the local animal committee (School of Chemistry, UNC, Exp. 0007526/2018). Male albino Wistar rats (12-15 weeks), inbreed in our laboratory for 5 years, were maintained on 12:12 h light-dark cycles with lights on (less than 50 lux of white fluorescent lamp) from zeitgeber time (ZT) 0 to 12 from birth until the day of the experiment. Food and water were available *ad libitum*.

### Retinal Light Damage

Retinal degeneration was induced as described by Contín et al. [6]. Briefly, animals were exposed to constant light in boxes equipped with LED devices (EVERLIGHT Electronic Co., Ltd. T-13/4 3294-15/T2C9-1HMB, colour temperature of 5,500 K) in the inner upper surface and temperature-controlled at 24 ± 1°C. At rat eye level, 200 lux were measured with a light meter (model 401036; Extech Instruments Corp., Waltham, MA, USA). After light stimulation the animals were sacrificed in a CO2 chamber at ZT6.

### Light Exposure Protocol

Animals were exposed to constant light stimulation for 2, 4, 6, and 8 days (LL2–LL8). Rats exposed to fluorescent light at 50 lux on 12:12 h light-dark cycles (LDR) or in constant darkness (DD7), were used as controls of standard housing conditions.

### Immunohistochemistry

After exposure, whole rat eyes were fixed in 4% (w/v) paraformaldehyde (PFA) in phosphate-buffered saline (PBS, pH 7.3) overnight at 4°C, cryoprotected in sucrose and mounted in optimal cutting temperature compound (OCT; Tissue-Tek^®^ Sakura). 20-μm-thick retinal sections were obtained along the horizontal meridian (nasal-temporal) using a cryostat (HM525 NX-Thermo Scientific). Sections were washed in PBS and permeabilized with PBS 0.2% (v/v) Triton X-100, 40 min at room temperature (RT). Then, they were blocked with blocking buffer (PBS supplemented with 0,05% Triton X-100; 3% (w/v) BSA, 2 % (W/V) horse serum and 0.2 % (w/v) Sodium Azide) for 2:30 h at RT with continuous gentle shaking. After that sections were incubated with rabbit polyclonal anti-glial fibrillary acidic protein (GFAP, Cat. No.G9269, Sigma–Aldrich Co., St. Louis, MO, USA, dilution 1:500), Goat polyclonal anti-ionized calcium binding adaptor molecule 1 (Iba1, Cat. No. ab107159, Abcam, Cambridge, UK, dilution 1:500) diluted in a blocking buffer, overnight (ON) at 4°C in a humidified chamber. Samples were then rinsed three times by 5 min in PBS 0.05% (v/v) Triton X-100 and incubated with goat anti-rabbit 549 IgG and Donkey anti goat 488, respectively and 3 μM DAPI, for 1 h at RT. Finally, they were washed 3 times in PBS and mounted in Mowiol (Sigma-Aldrich). Images were collected using a confocal microscope (Olympus FV1200, Japan).

### Microglia cells number analysis

To analyze the number of microglial cells, vertical cryosections of retina immunostained with Iba1 were used. The quantification of the number of Iba1 positive cells were defined in two regions: a) *inner portion retina*, corresponding to ganglion cell layer (GCL) and IPL, inner nuclear layer (INL) and OPL and b) *outer portion retina*, corresponding to ONL + outer segment (rods and cones). Three non-consecutive sections, taken in fields on both sides of the optic nerve area per animal from each group, were analyzed (n=3 retinas/group).

### Western Blotting

Homogenates of the whole retina suspended in a 200 μl PBS containing protease inhibitors, were lysed by repeated cycles of ultra-sonication, and total protein content was determined by the Bradford method [24]. Lysates were then suspended in sample buffer [62.5 mM Tris–HCl pH 6.8; 2% (w/v); SDS, 10% (v/v); glycerol; 50 mM DTT; 0.1% (w/v) bromophenol blue] and heated at 90°C for 5 min. Proteins (25 μg/lane) and molecular weight marker [5 μl ECL Rainbow Marker-Full range (12,000–225,000 Da) from Amersham Code RPN7800E] were separated by SDS-gel electrophoresis on 10% polyacrylamide gels, transferred onto nitrocellulose membranes, blocked for 1 h at RT with blocking buffer consisting of 5% (w/v) skim milk in washing buffer [(0.1% (v/v) Tween-20 in PBS], and then incubated overnight at 4°C with anti-GFAP, or anti-α-Tubulin antibody diluted 1:1000 in blocking buffer. The membranes were subsequently washed, and incubated with the corresponding secondary antibody (Goat anti-rabbit IRDye ^®^ 700CW or Goat anti-mouse IRDye ^®^ 800cW, Odyssey LI-COR) in PBS for 1 h at RT, followed by three washes (5 min each wash) with washing buffer. Membranes were scanned using an Odyssey IR Imager (LI-COR Biosciences) and the quantification of the protein bands was performed by densitometry using the FIJI / Image J program (NIH).

### RNA Extraction, cDNA synthesis and real time RT-PCR

Total RNA was extracted from individual rat retinas using TRIzol^®^ RNA extraction reagent (Invitrogen, Carlsbad, CA), according to the manufacturer’s instructions. The isolated RNA was quantified using an Epoch Microplate Reader (BioTek Instruments, Winooski, VT, USA). For complementary DNA (cDNA) synthesis, 2 μg of RNA was treated with DNase I (Thermo scientific, USA) to remove possible contamination with genomic DNA. The product was incubated with a mix of random hexamer and Oligo-dT primers (Biodynamics), deoxynucleotides and the reverse transcriptase M-MLV (Promega, Madison, WI, USA), in RNAse-free conditions. Reverse transcription was performed following the manufacturer’s specifications, employing a thermocycler Mastercycler gradient (Eppendorf, Hamburg, Germany). Real time RT-PCR was performed using the CFX96 Touch Real-Time PCR Detection System (Bio-Rad, USA). cDNA (120 ng) was amplified in 15 μl reaction mixture consisting of 7.5 ul of 2x SYBR Green PCR Master Mix (Life Technologies, USA), 0,75 μl 10 μM primer mixture and 0,75 μl of Nuclease-Free Water. The parameters used for PCR were as follows: 95°C for 5 min (1 time); 95°C for 30 sec, 60°C for 30 sec, and 72°C for 30 sec (40 times); and 95°C for 60 sec (1 time). Samples were subjected to a melting-curve analysis to confirm the amplification specificity. Semi-quantification was performed by the method of ΔΔCt. The fold change in each target gene relative to the □-Actin endogenous control gene was determined by: fold change = 2^-Δ(ΔCt)^ where ΔCt = Ct_(target)_ - Ct_(□-Actin)_ and Δ(ΔCt) = ΔCt_(LL)_ - ΔCt_(LDR)_. real time RT-PCR were run separately for each animal in triplicate. The primers used for real time RT-PCR are provided in Table 1.

**Table 1:**
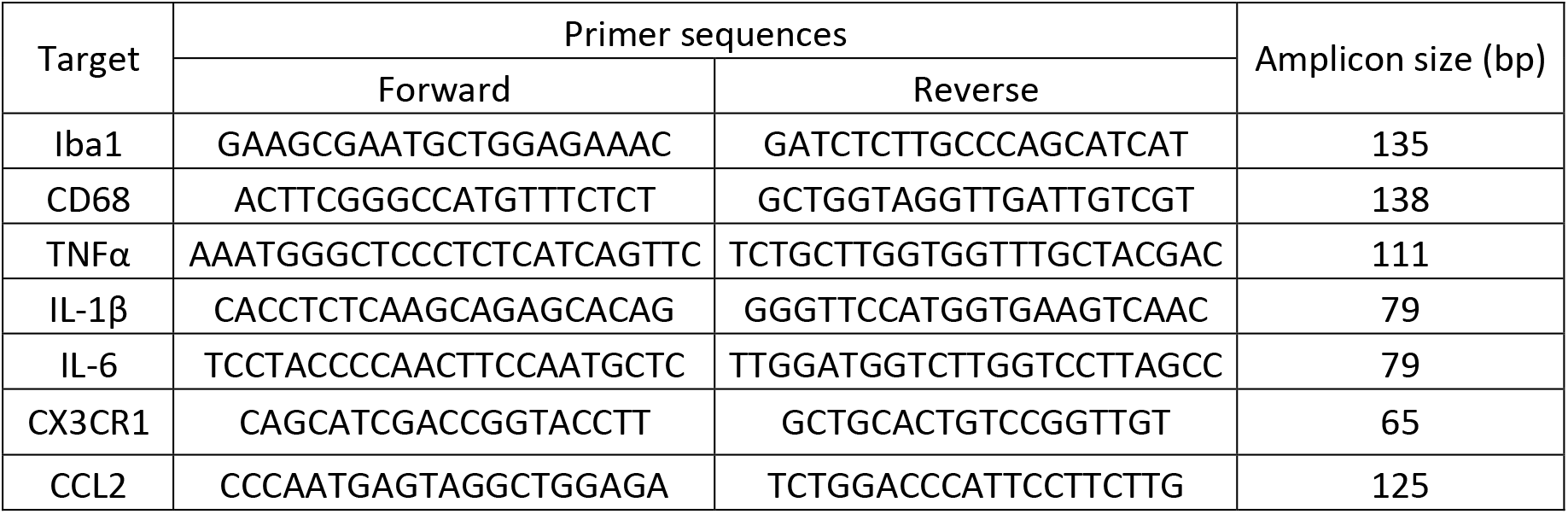
Primer sequences of the genes related to glial cell markers, inflammatory and chemokines.

## STATISTICAL ANALYSIS

Statistical analysis was carried out using the Infostat software (Version 2017, InfoStat Group, FCA, National University of Córdoba, Argentina). The normality and homogeneity of the variance assumptions were proved with Shapiro-Wilks and Levene tests, respectively. Data were analyzed using one-way analysis of variance (ANOVA) and Tukey’s *post hoc* test. A non-parametric Kruskal-Wallis test was performed when the data did not comply with the assumptions of the ANOVA. Data are expressed as mean ± SEM. In all cases, a *p* value < 0.05 was considered statistically significant. All graphics were made using GraphPad Prism Software, version 6.01 (San Diego, CA, USA).

## RESULTS

As we demonstrated before, treatment of rat retinas with LED light for constant several days resulted in a diminution of ONL corresponding to photoreceptor cells (Fig. 1a, LL6) [6]. Thus, along with the degeneration of the outer retina, as part of the mechanism of light damage, the immune response may be triggered possibly by the activation of retinal glial cells. Thus,we characterized the macro and microglia response, in order to evaluate the glia activation during the onset of RD in Wistar rats induced by constant exposure to 200 lux of LED light.

**Figure 1.**
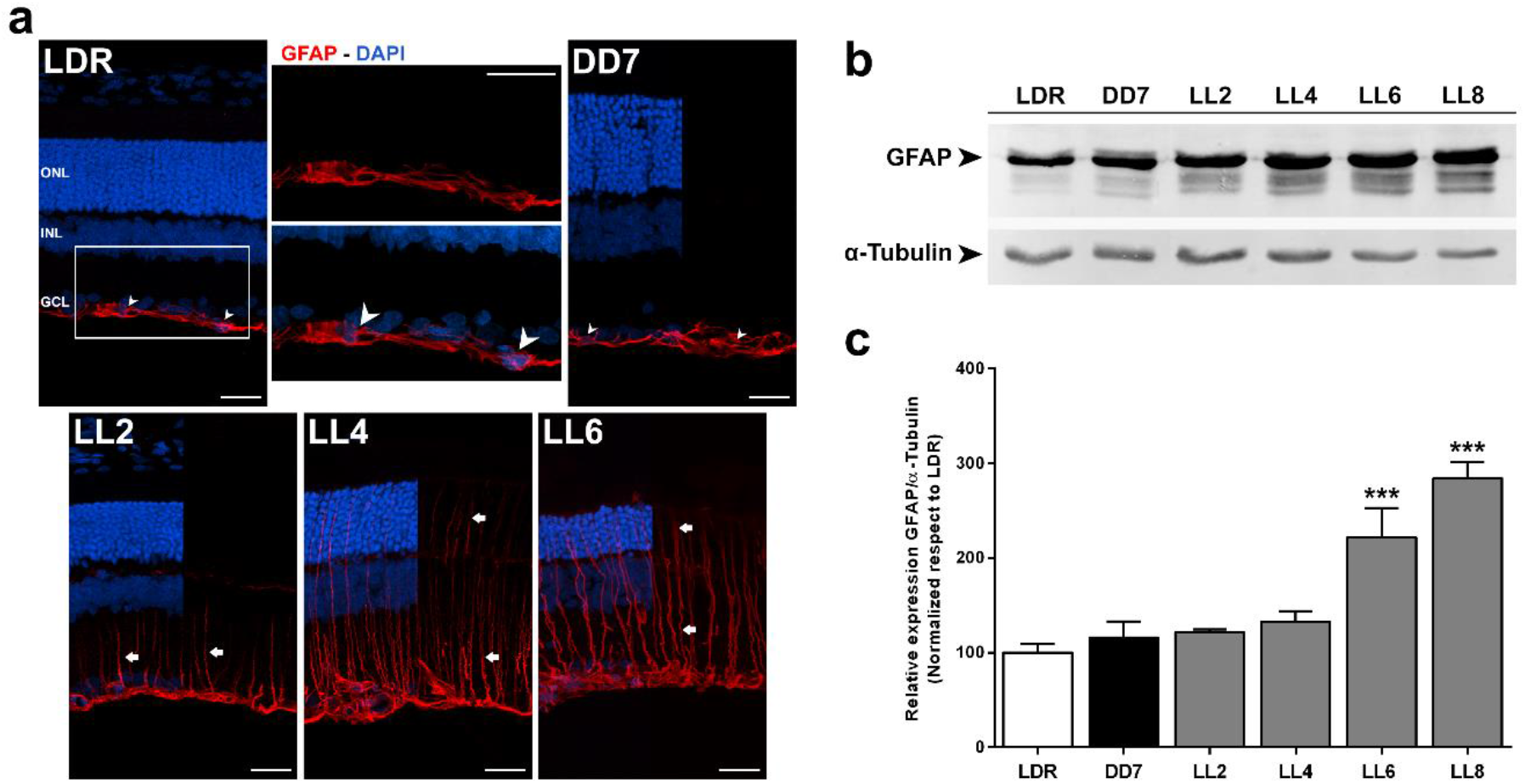
Effect of constant light exposure on GFAP expression in rat retinas. **a.** Immunofluorescence staining showing GFAP expression in rat’s retinas exposed to light (LL), constant darkness (DD7) and light-dark cycles 12-12hs (LDR); red: GFAP antibody staining; blue: nuclear DAPI staining. Magnification 40x (insert 60x). Scale bar = 25 μm. ONL, outer nuclear layer; INL, inner nuclear layer; GCL, ganglion cell layer. **b.** Analysis of GFAP expression by Western Blot, a representative werten blot of GFAP and α-Tubulin staining is showing, BDPs, Breakdown products. **c.** Densitometric quantification of wester blot staining with GFAP and α-Tubulin is shown as relative expression (GFAP/α-Tubulin) expressed as percentage and normalized respect to LDR. Data are presented as mean ± SEM. \*\*\**p* < 0.001, \*\**p* < 0.01, \**p* < 0.05 by ANOVA and Tuckey’s *post hoc* test. n=4 animals/group.

### Macroglia

#### Immunohistochemistry and western blot

In healthy retina, GFAP is expressed in Müller cells end feet but in the presence of metabolic stress, it is upregulated and expressed in the entirety of these cells, becoming an important marker of reactive gliosis [25]. Also, GFAP is a typical marker for retinal astrocytes, present in reactive and non-reactive cells; in retina, GFAP immunostaining labels astrocytes in the GCL and their processes in the nerve fiber layer, and in blood vessels of the superficial plexus [26]. After an injury, both macroglial cells have been implicated in retinal gliosis. Müller cell gliosis effects are an enigma of major retinal diseases, [10]; Hence, in order to study the retinal macroglia response during retinal injury promoted by light, we analyzed GFAP expression in control retinas and after different times of light exposure (LL).

Immunostaining for GFAP in retina from control groups LDR and DD7 showed intense labeling in GCL and short filament projections (Figure 1a, white arrowhead); this morphology and localization of GFAP positive cells was indicative of retinal astrocytes. However, after LL2, the expression of GFAP increased, and GFAP labeling extended into an elaborate filamentous structure spanning the retinal thickness, indicative of Muller cells expression, with higher levels at LL6 (Figure 1a, white rows). Western blot analysis (Figure 1b) showed an increase of GFAP and breakdown products (BDPs) at LL2, LL4, LL6 and LL8 respective to LDR. Data quantification demonstrated a gradual increase of GFAP expression along with time of exposure, reaching a statistically significant increase at LL6 and LL8, compared to control groups (LDR and DD7) and experimental groups, LL2 and LL4 (Figure 1c).

### Microglia

#### Immunohistochemistry and cell number analysis

In order to assess microglial cell activation in retinal light damage promoted by exposure to constant low light, we study the shape and localization of these cells by immunolabeling of retinal sections with Iba1 antibody. Iba1 is a microglia marker widely used for studies of microglia in resting or activated states. The morphology of this cell is strongly related to the activation; it is accepted that resting inactive microglia have an extremely dynamic morphology with a small cell body and many long and ramified processes. When microglia become activated, the cells display morphological changes such as size increase, retraction and thickening of the processes, and deformation of the cell soma acquiring an “ameboid shape” [27].

After immunostaining with Iba1, we found few and weak-labeled positive cells in GCL, IPL and OPL in the control retina (LDR and DD7), and these cells showed typical resting microglia characteristics: small cell body with long and ramified processes (Figure 2a). In retinas exposed to light treatment, we found an increasing number of Iba1 positive cells in INL, IPL and the labeling in OPL and ONL, at LL4 and LL6. The cell morphology became ameboid with thicker processes characteristic of activated cells (Figure 2a). Collectively, these data suggested that Iba1 positive cells, macrophages, are activated, increasing in number and accumulation in the outer retina in our model of RD promoted by light.

**Figure 2.**
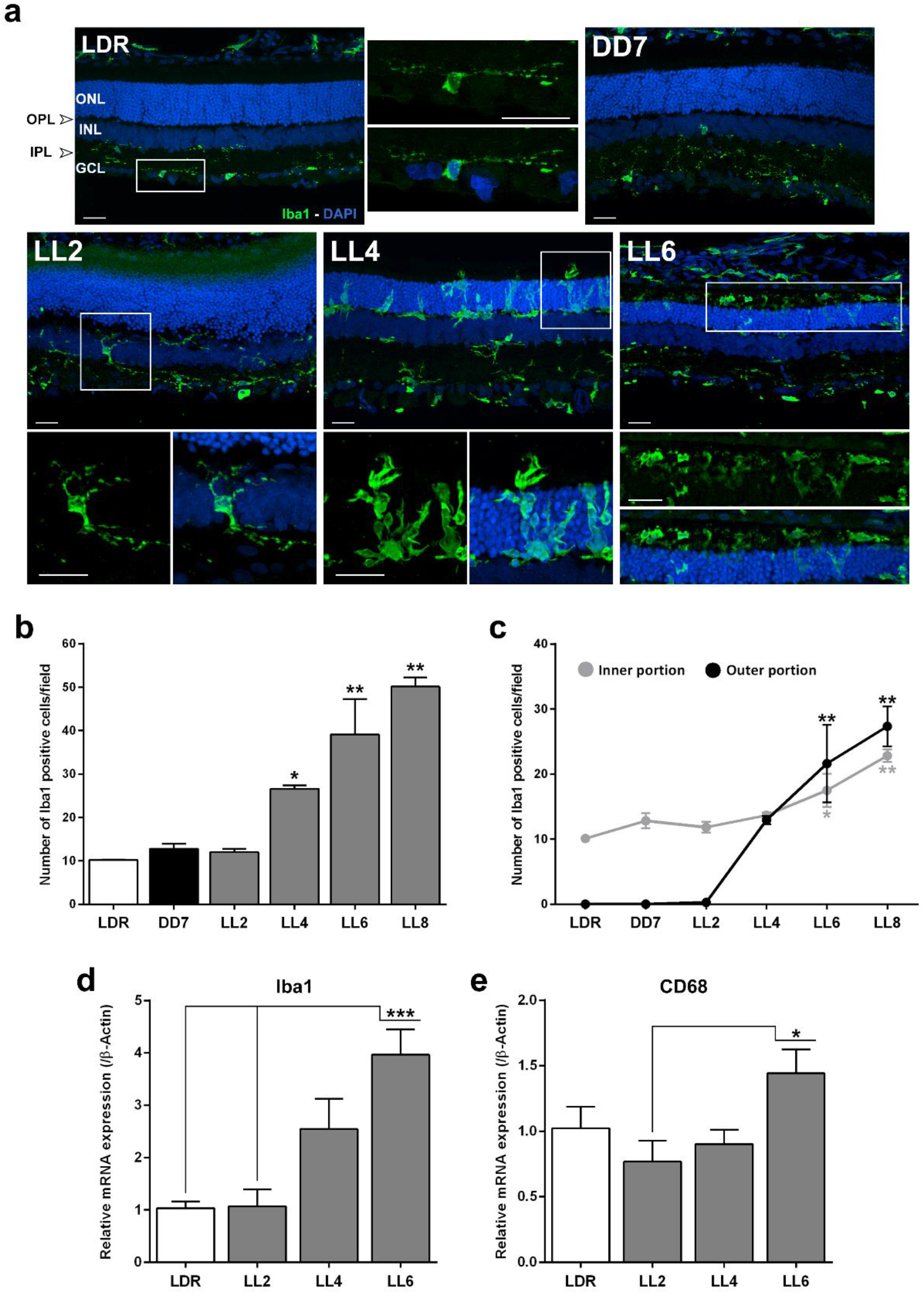
Microglial cell response as a consequence of constant light stimulation. **a.** Iba1 immunostaining in animals exposed 2, 4 and 6 days to light (LL2-6) and in control groups (DD7, LDR). Green: Iba1 antibody staining, blue: nuclear DAPI staining. Magnification 40x (inserts 60x) Scale bar = 25 μm. ONL, outer nuclear layer; INL, inner nuclear layer; GCL, ganglion cell layer; IPL, inner plexiform layer; OPL, outer plexiform layer. **b.** Iba1 positive cells quantification in central retina. Three non-consecutive sections of one retina per animal were analyzed (spaced at least by 50 μm). Data are presented as mean ± SEM. \**p* < 0.05, \*\**p* < 0.01 by Kruskal-Wallis test. n=3 animals/group. **c.** Comparison of the number of Iba1 positive cells in *inner portion* (GCL, IPL, INL + OPL) and *outer portion* (ONL + OS). Data are presented as mean ± SEM. \**p* < 0.05, \*\**p* < 0.01 respect to LDR, ^#^*p* < 0,05 respect to DD7 and LL2 by Kruskal-Wallis test. n = 3 animals/group. **d.** Quantitative analysis of Iba1 and CD68 mRNA by real time RT-PCR. Data are presented as mean ± SEM. \**p* < 0.05, \*\*\**p* < 0.001 by one way ANOVA and Tuckey’s *post hoc* test. n=5 to 7 animals/group.

In order to further the analysis of microglial response in retinas exposed to constant light, we counted Iba1 positive cells in two retinal areas: a) *inner portion retina*, corresponding to GCL and IPL, INL and OPL and b) *outer portion retina*, corresponding to ONL + outer segment (rods and cones) (Figure 2c). When we analyzed the two areas defined, we found a higher number of Iba1 positive cells in the *outer retina* with a statistical significance after LL6 respect to controls (Figure 2c), indicative of profuse migration and activation of these cells after LL6. The analysis of total Iba1 positive cells along all the cell layers confirmed this result showing a significant increase of Iba1 positive cells after LL4 and LL6 respective to control animals (Figure 2b).

#### Relative expression levels of Iba1

In order to further confirm the increase in Iba1 expression, mRNA levels of Iba1 were measured by real time RT-PCR. As shown in Figure 2d, while mRNA levels of Iba1 were similar in controls and LL2 retinas, the expression increased 2- and 4-fold at LL4 and LL6, respectively. Statistical analysis showed significant upregulation of Iba1 transcript levels at LL6 respect to LDR (*p* < 0.05).

Altogether, our results indicate that low light promotes retinal microglial activation after LL6 of constant exposure.

#### CD68 expression

Several surface markers such as CD68, complement receptor 3 (CD11b/CD18, OX42), MHC-II (OX6), F4/80, and Griffonia simplicifolia isolectin B4, are also used to detect and classify microglia activity [28]. In order to complete the characterization of microglia response, we studied the expression of CD68 by real time RT-PCR. CD68 is a glycoprotein localized in lysosomes and a marker for activated phagocytic cells [29]. CD68 expression (Figure 2e) did not change after light exposure at LL2 and LL4 relative to control LDR, while it increased significantly at LL6 (*p* < 0.05), supporting the idea that the microglial activation reaches significant levels after 4 to 6 days of exposure.

#### Inflammatory mediators

Activated microglia may exert detrimental neurotoxic effects by excessive production of cytotoxic factors such as tumor necrosis factor (TNF) α, interleukin (IL)-1β, IL-6, reactive oxygen species, among others. The cytotoxic factors may amplify the cascade of microglial activation, causing neurodegeneration [30, 31]. Therefore, we studied the expression levels of immune mediators by real time RT PCR. As shown in Figure 3, relative mRNA levels of TNFα (Figure 3a; *p* < 0.5) and IL-6 (Figure 3c; *p* < 0.5) were significantly increased only at LL6 compared with control LDR. IL-1β did not show significant changes of mRNA expression at any of the times studied (Figure 3b). All these results suggest that inflammatory mediators are induced by constant light after six days of exposure.

**Figure 3.**
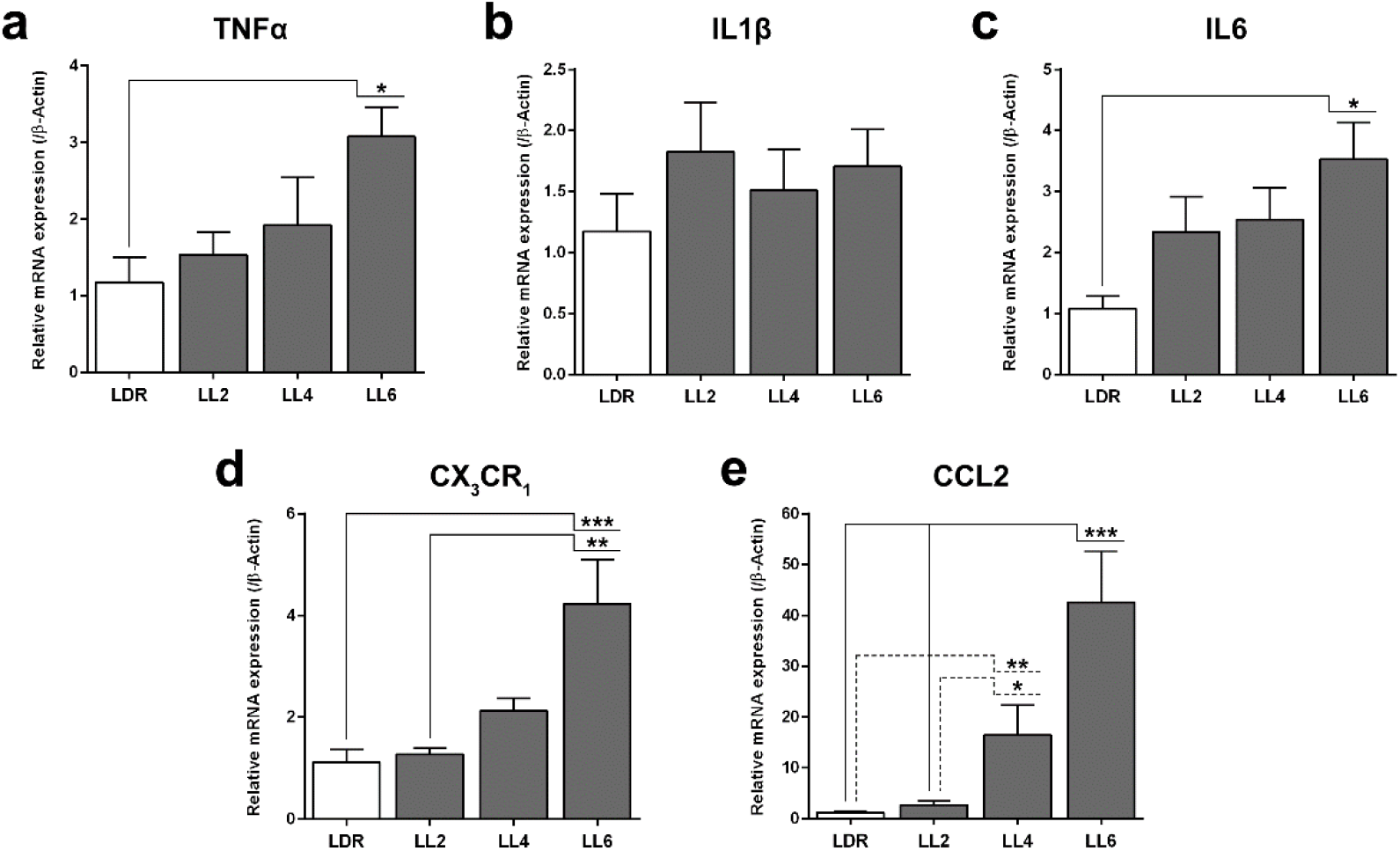
Quantitative analysis of pro-inflammatory markers at the level of mRNA expression by real time RT-PCR after light exposure. **a.** TNFα, **b.** IL-1β, **c.** IL-6, **d.** CX3CR1 and **e.** CCL2. In all cases, data are presented as mean ± SEM. \**p* < 0.05, \*\**p* < 0.01, \*\*\**p* < 0.001 by one way ANOVA and Tuckey’s*post hoc* test. n=5 to 7 animals/group.

Chemokines are a family of small signaling proteins that regulate monocyte/macrophage activation and recruitment, acting as chemoattractant and activating inflammatory cells [32]. Chemokines are involved in the pathogenesis of immune-mediated inflammation [33] and their levels increased significantly after inflammation-associated neurodegeneration. Fractalkine is a chemokine belonging to the CX3C subfamily that exists as both soluble and membrane bound forms [34] and it has been found in retinas exposed to bright light damage [35]. In AMD progression, the major chemokine signaling pathways linked are CCL2/CCR2 and CX3CL1/CX3CR1 [36]. Chronic stress can also stimulate the expression of CCL2, increasing monocyte and microglial cell recruitment and amplifying the inflammatory state [37]. In order to evaluate fractalkine and CCL2 involvement in low light induced-damage, we studied the fractalkine receptor (CX3CR1) and CCL2 mRNA expression during continuous light exposure. We found that both CCL2 and CX3CR1 expression increased significantly from LL4 and LL6 respectively, in comparison to control retina (LDR); (Figure 3d and e).

## DISCUSSION

The excess of artificial light is a growing problem called Light pollution. It produces several complications at different ecological and environmental levels; in vision, it might produce retinal degeneration *per se* or accelerate other retinal diseases [1]. The possibility to understand the molecular mechanisms of light damage in different retinal injury models will contribute to the knowledge of visual disorders related to light pollution and defects in the phototransduction mechanism. In the present study, a Wistar rat model of constant light-induced RD was used to study the time-course of glial activation. As we demonstrated before, we found a reduction in ONL of rat retinas after LL6 compared with control animals LDR and DD7 (Figure 1). GFAP immunostaining analysis showed differences between control and LL treated animals, demonstrating an increase in the number of GFAP-positive cells with a filamentous morphology through the retinal thickness which extend to ONL (Figure 1a, white arrows). GFAP is an important marker of reactive gliosis, that is up-regulated and expressed in the entirety of Müller and astrocytes cells in the presence of stress [52, 53]. Therefore, our results confirm a clear activation of macroglia after constant exposure to low light (Figure 1). Analysis of GFAP expression by western blot showed a significant increment of GFAP and BDPs levels at LL6 and LL8, respective to control LDR (Figure 1c). It has been shown that the proteolytic conversion of intact glial protein GFAP (50 kDa) into BDPs is involved in glial injury in CNS during the acute/subacute phase of several neurotrauma, and GFAP-BDPs were associated with the role of Ca^+2^-dependent protease calpains or cathepsin D as alternative death pathways [25]. Previously, we demonstrated that a caspase-3 independent cell death mechanism is involved in retinal damage induced by constant low light exposure, suggesting the participation of other ways of cell death [6]. Ca+2-dependent protease calpains or cathepsin D as alternative death pathways may involve a late gliosis mechanism, in which GFAP-positive cells may exert cytotoxic effects on the retina after LL6; this mechanism has been described for other retinal light damage [38–41].

Immunohistochemical analysis of the retina showed Iba1 positive cells in the IPL and OPL in the control retinas (LDR and DD7); the characteristic shape of the cell as small cell body and long and ramified processes indicates resting microglia cells (Figure 2a); however, at LL4, LL6 and LL8, increasing levels of Iba1 positive cells in INL and IPL were observed, as well as new Iba1 positive cells in OPL and ONL with a characteristic ameboid morphology and thick processes, revealing macrophages and microglial activation promoted by light. The quantification of Iba1 positive cells in two well-defined zones called *inner portion retina* and *outer portion retina* (Figure 2c) showed a significant increase in cell numbers in the *outer portion retina* after LL4 and LL6, respective to controls, indicating profuse migration and activation of Iba1 positive cells after LL6. Analysis of total Iba1 positive cells, showed a significant increase from LL4 respect to controls animals (significant value at LL4, LL6 and LL8 respect to LDR with a *P* = 0.0389, *P* = 0.0029 and *P* = 0.0013, respectively) (Figure 2b), demonstrating important microglial activation after LL4. mRNA expression confirms these results showing that Iba1 levels increased at LL4 to LL8 (significant value at LL6 respect to LDR with a *P* < 0.001) while no changes were observed in controls and LL2 animals (Figure 2d). Similarly, CD68 mRNA analysis showed significantly increased levels of this marker at LL6 (*P* < 0.05), supporting the fact that constant exposure to low light induced a late microglial activation (Figure 2e).

As retina is the tissue adapted to capture light photons, it is continuously exposed to light and oxygen, making it vulnerable to oxidative stress [1]. Therefore, the continuous surveillance of the retina for the detection of noxious stimuli is mostly carried out by microglial cells, the resident tissue macrophages which confer neuroprotection against transient pathophysiological insults. Under sustained injury, like constant light, microglial inflammatory responses might become deregulated, promoting photoreceptor cell death via other mechanisms. In this sense, oxidative stress is an important factor in injured retina frequently involved in worsening disease progression, which is induced by events such as hypoxia or inherited mutations, and triggers microglial activation [21, 42, 43]. Previously, we demonstrated the existence of oxidative reactions in LL exposed rats [7]. Dihydroethidium (DHE)-positive labeling in the retina ONL, and a significant increase in ROS production aty LL5, were both indicative of active oxidative stress responses after 5 days of constant light. Thus, oxidative stress may be related to glial activation after several days of constant light. During the first days of light exposure, microglia may enhance tissue repair processes in order to return to homeostasis; however, under sustained light stimulus, the induction of microglial inflammatory responses may promote photoreceptor cells death. Our data showed an upregulation of pro-inflammatory factors TNFα and IL-6; relative mRNA levels increased significantly at LL6 compared with control LDR (Figure 3a and c). However, IL-1β did not show significant changes of mRNA expression at any times of LL studied (Figure 3b). It is known that upon injury, microglia and monocytes are activated promoting the secretion of neurotoxic factors, such as interleukin IL-1β and TNFα, contributing to neurodegeneration [36,44]. In bright light-induced photoreceptor degeneration models, retinal microglia and monocytes are activated and recruited to the outer retina area where photoreceptor apoptosis occurred; this activation was correlated with the upregulation of the proinflammatory factors IL-1β and TNFα [45]. Our time-course study in LL confirmed that low light exposure promotes the upregulation of the proinflammatory TNFα and IL-6 in a time-related manner, in parallel with microglial activation and migration; nonetheless this activation is late in LL, after the initial photoreceptor cell death occurred, indicating that the inflammatory responses may be part of a secondary mechanism secondary to an initial signal cascade yet to be elucidated. In bright light damage, it has been shown that the apoptotic peak of photoreceptors was consistent with an elevated level of fractalkine, and both alterations were ahead of the peak of microglial infiltration; these observations suggest that after exposure to intense blue light, soluble fractalkine is initially released by injured photoreceptors, and thereby causes the migration of microglia into the ONL via CX3CR1 [35]. Here we demonstrated that in low light damage both CX3CR1 and CCL2 mRNA expression increased after LL6 supporting a role for these chemokines at the time of active gliosis and inflammation, and suggesting an ongoing cross-talk between microglia and neuronal cells at later stages of neurodegeneration (Figure 3 d and e).

Studies in Arr1-/- (visual arrestin, intracellular protein that desensitize rhodopsin) retina mouse model showed that microglia and monocyte proliferation induced photoreceptor degeneration, and CCL2-CCR2 pathway was identified as an important mediator of retinal health; however, Microglia and monocyte proliferation occur concurrently, but not until several days after photoreceptor [53], supporting the idea that other inicial mechanism of photoreceptors cell death. In photoreceptors lacking Arrestin-1, rod phototransduction pathway is greatly affected since it prolongs rhodopsin activation; this effect results in light-induced photoreceptor degeneration and it isone of the mains triggers of cell death [46, 47]. We think that the late macro and microglia activation may be involved in retinal degeneration in low light retinal damage; however, some phototransduction-dependent factors may be the initial event that set off death pathways cascades. Electroretinograms studies performed in animals under constant light exposure (LL), we previously demonstrated that both, “a” and “b” waves tend to decrease the amplitudes and increase the latency time during light stimuli, getting abolished records before LL4 [48]; these results show that electrical dysfunction activity precedes ONL cell loss, redox imbalance [7], and macroglial (Figure 1,) and microglial (Figure 2,3) activation, indicating that the initial death signal occurred before LL4. Low intensity light stimuli need the activation of photopigments and phototransduction mechanisms to induce retinal degeneration [2, 49], suggesting impairment of the phototransduction mechanism could be responsible for cell death. Moreover, we previously demonstrated the existence of more phosphorylated rhodopsin (rhodopsin phospho-Ser334) after LL2 with a higher level at LL7, supporting the idea that early changes in phototransduction cascade are involved [6]. Further studies are necessary to determine the role of opsin-mediated retinal degeneration processes in this model; however, we think that retinal dysfunction during the first days of LL (LL1-LL4) may be promoted by phototransduction processes deregulation which may induce other cell death pathways. It is important to highlight that the maximal glia activation and proinflammatory responses concurred at LL6, later than the onset of photoreceptor cell death, indicating that other mechanisms may be initially involved but are exacerbated by the action of gliosis. We speculate that one of the key mechanisms for the initiation of this process could be rhodopsin hyper-phosphorylation as a consequence of impairment of the enzymes related to the phosphorylation/dephosphorylation processes [6] being a phototransduction-dependent mechanism, according to observations previously described by Hao and collaborators [2].

In summary, using a RD model induced by chronic low intensity LED light exposure, we demonstrated that glial and immune activation appear to be secondary to others inicial neurodegeneration events and may be a phototransduction-dependent mechanism.

## FUNDING

This work has been supported by Consejo Nacional de Investigaciones Científicas y Tecnológicas de la República Argentina (CONICET), PICT 2013 No. 106 and CONICET PIP Nro. 11220150100226 (2015-2017) and (PIP) 2014-201. Secretaría de Ciencia y Tecnología—Universidad Nacional de Córdoba (SeCyT-UNC) 2014-2018 and 2018-2024, and by Agencia Nacional de Promoción Científica y Técnica FONCyT, PICT 2016 No. 0187 and FONCyT.

## ACKNOWLEDGMENTS

We are grateful to Dra. Cecilia Sampedro and Dr. Carlos R. Mas for technical support and Rosa Andrada for animal facility management.

## Notes

### Competing Interest Statement

The authors have declared no competing interest.

